# Induced electric fields inhibit breast cancer growth and metastasis by modulating the immune tumor microenvironment

**DOI:** 10.1101/2024.04.14.589256

**Authors:** Manish Charan, Travis H. Jones, Dinesh K. Ahirwar, Nandini Acharya, Vish V. Subramaniam, Ramesh K Ganju, Jonathan W Song

**Affiliations:** Department of Pathology, College of Medicine, The Ohio State University, Columbus, OH, 43210, USA; Department of Mechanical and Aerospace Engineering, College of Engineering, The Ohio State University, Columbus, OH, USA; Department of Bioscience & Bioengineering, Indian Institute of Technology, Jodhpur, India; Department of Neurology, College of Medicine, The Ohio State University, Columbus, OH, 43210, USA; Arthur G. James Comprehensive Cancer Center and Richard J. Solove Research Institute, The Ohio State University Medical Center, Columbus, OH 43210, USA; Pelotonia Institute for Immuno-Oncology, The Ohio State University Comprehensive Cancer Center – Arthur G. James Cancer Hospital and Richard J. Solove Research Institute, The Ohio State University, Columbus, OH 43210, USA

**Keywords:** Alternating electromagnetics, physics of oncology, immunomodulation, bioengineering

## Abstract

Despite tremendous advances in oncology, metastatic triple-negative breast cancer remains difficult to treat and manage with established therapies. Here, we show in mice with orthotopic triple-negative breast tumors that alternating (100 kHz), and low intensity (<1 mV/cm) induced electric fields (iEFs) significantly reduced primary tumor growth and distant lung metastases. Non-contact iEF treatment can be delivered safely and non-invasively *in vivo* via a hollow, rectangular solenoid coil. We discovered that iEF treatment enhances anti-tumor immune responses at both the primary breast and secondary lung sites. In addition, iEF reduces immunosuppressive TME by reducing effector CD8+ T cell exhaustion and the infiltration of immunosuppressive immune cells. Furthermore, iEF treatment reduced lung metastasis by increasing CD8+ T cells and reducing immunosuppressive Gr1+ neutrophils in the lung microenvironment. We also observed that iEFs reduced the metastatic potential of cancer cells by inhibiting epithelial-to-mesenchymal transition. By introducing a non-invasive and non-toxic electrotherapeutic for inhibiting metastatic outgrowth and enhancing anti-tumor immune response *in vivo,* treatment with iEF technology could add to a paradigm-shifting strategy for cancer therapy.

## Introduction

Breast cancer is the most frequent malignancy in women and the leading cause of death in women in many countries^1^. Triple-negative breast cancer (TNBC) represents 10-15% of all breast cancers reported^2^ and is the subtype that is the most challenging to treat given its aggressive phenotype and limited molecular targets for therapy^3^. Even if diagnosed early, TNBC has a high risk for relapse and mortality^4^. Neoadjuvant therapy for TNBC still relies heavily on conventional cytotoxic chemotherapy drugs^5^. Yet, chemotherapies are frequently associated with adverse events that compromise quality of life and lead to early treatment discontinuation^6^. Current treatment options with manageable toxicity profiles for TNBC patients are therefore limited and frequently ineffective. This unmet need underscores the urgency for new, effective, and well-tolerated therapeutic approaches for TNBC.

Breast cancer, especially during the conversion from localized to metastatic disease, is characterized by a tumor microenvironment (TME) of progressive immune suppression^7^. This is evident by the recruitment of cell types that suppress anti-tumor immune reactivity (i.e., are immunosuppressive) such as M2-like tumor associated macrophages (TAMs), myeloid-derived suppressor cells (MDSCs), and T regulatory cells (Tregs)^8,9^. The recruitment of these immunosuppressive cells reduces the effector function of CD8+ T cells^10–12^. This immune landscape has led some to describe breast cancer as immunologically “cold”^13^. Thus, interventions that target different immunosuppressive elements within the TME may expand clinical responses to immunotherapy in metastatic breast cancer, including TNBC.

Low-intensity induced electric fields (iEFs) produced by exogenously applied temporally varying magnetic fields offer great promise in treating cancer in a non-invasive, non-contact, and non-pharmacological manner^14–16^. It has been previously shown that iEFs act selectively on human TNBC cell lines (MDA-MB-231 and MCF10CA1a) compared to normal epithelial breast cells (MCF10A)^14–16^. These selective effects of iEF treatment include hindering cancer cell migration in the presence of chemokines and growth factors^14–16^, rendering their actin cytoskeleton diffuse and unable to form filopodia, and improving the efficacy of Akt inhibitor MK2206 in hindering their migration^14^. iEF treatment also significantly reduced the activity of succinate dehydrogenase (SDH) in the mitochondria of cancer cells, thereby interfering with their ability to produce ATP via oxidative phosphorylation^15^. While iEFs have demonstrated direct and selective tumor suppressive signaling responses in cancer cells *in vitro*, their effects in the context of an *in vivo* TME with an intact immune system are unknown.

Here, we describe the first *in vivo* application of iEFs in a mouse model of TNBC. We adapt a previously described non-contact method of applying iEFs *in vitro*^14^ for *in vivo* studies by inserting a mouse cage inside a Plexiglas open-ended box with a single-turn coil wrapped around the box. We demonstrate that iEF treatment inhibits primary tumor growth and distant metastases to the lung *in vivo*. We further show that iEF treatment enhances the anti-tumor immune responses in TNBC by reducing the immunosuppressive TME while demonstrating no adverse effects in treated mice. These results suggest that iEF treatment is a novel intervention for shaping an effective immune response against TNBC.

## Results

### Induced electric fields (iEFs) are safe and show no adverse events in mice

To investigate the *in vivo* effects of iEF treatment, we have devised an apparatus for controlled application of iEFs to mice in cages (**Fig. 1A-C**). The iEF apparatus comprises an open-ended, transparent, acrylic box, around which a single-layer coil is wound (**Fig 1A**, see Methods for coil characteristics). A 20 V_pp_, sawtooth-shaped, 100 kHz waveform was applied to the coil using a function generator (Hewlett Packard 33120A) to produce an iEF throughout the interior of the mouse cage. In this configuration, all mice present within the cage receive simultaneous, systemic treatment. **Figure 1B** shows the contours of calculated iEF in the y-z plane (long side of the box/cage) with magnitudes on the order of 1 mV/cm. In comparison, these iEF magnitudes are about 1-2 orders of magnitude larger than the maximum iEF reported in previous *in vitro* experiments^15–17^, and 3 orders of magnitude *smaller* than the ∼1 V/cm EFs reported with tumor treating fields (TTFs)^18,19^. As a further benchmark, the static electric field at the surface of the earth is on the order of 1 V/cm^20^. The non-negligible components of the iEF inside the box are in the x and y directions, with the iEF relevant to the location of the mice being primarily in the x direction toward the center of the cage and comparable in magnitude in both x and y directions near the edges of the cage floor (**Fig. 1B**). In the region of the cage occupied by mice, the calculated time-averaged values of the magnitude of iEF range from 0.67 mV/cm to 1.21 mV/cm with a spatial average value of 0.97 mV/cm (**Fig. 1C**).

**Figure 1:**
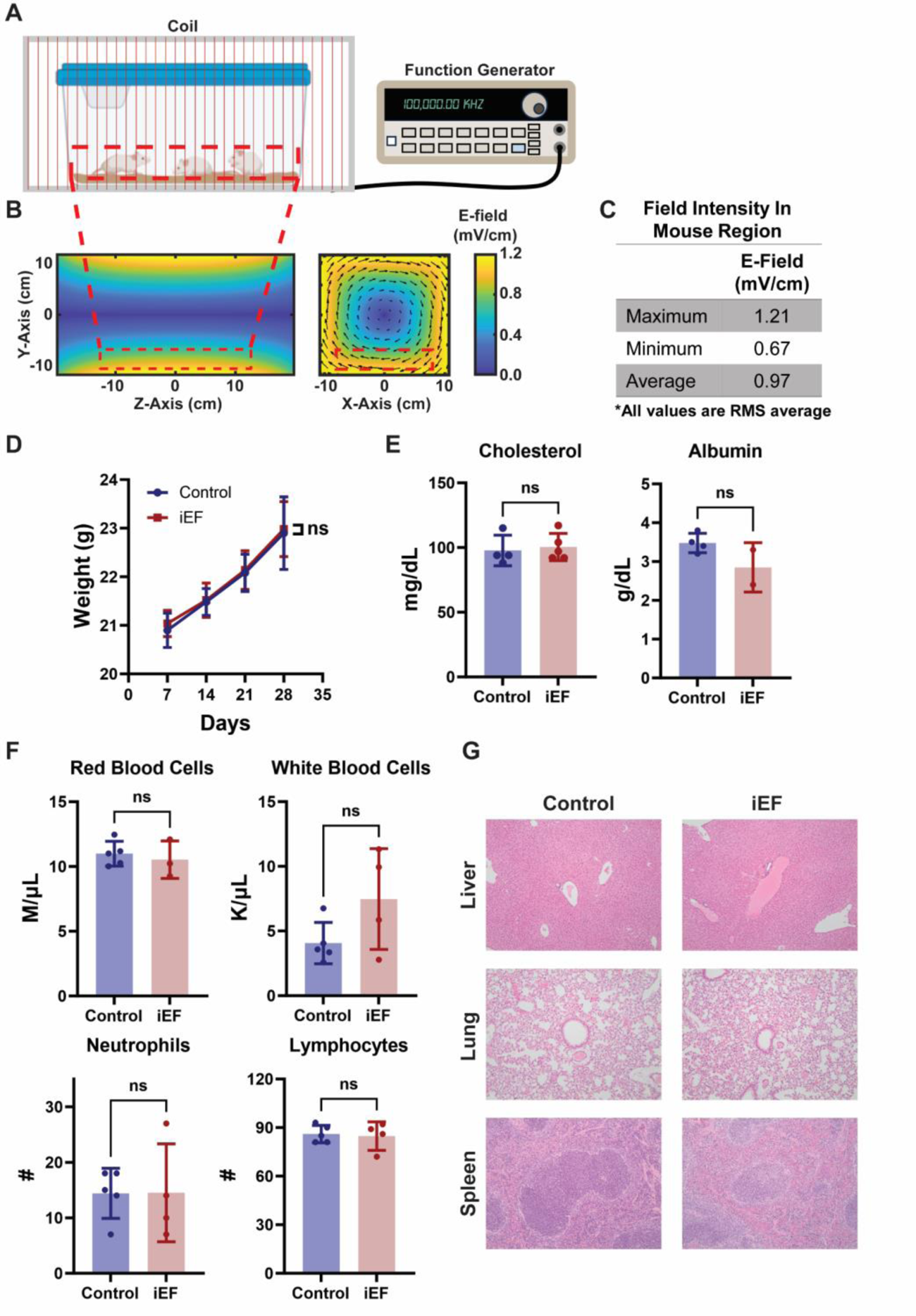
iEF treatment is well tolerated and safe in mice. **A)** Schematic of experimental setup. A function generator powers the electromagnetic coil with the plastic mouse cage sitting in the bore of the coil. Mice are free to move around within the cage. **B)** Contour plot of side- and front-view of electric field generated by coil. Black boundary represents the coil edges while the red, dashed boundary represents the region the mice primarily move around. Black arrows in front-view indicate direction of induced electric field during peak field strength. **C)** A summary of the electric field characteristics within the mouse region. Values are root mean square (RMS) time averaged. **D)** BALB/c mice were treated under iEF for 30 days. Body weight was measured every seven days. **E)** Blood and serum were extracted at endpoint for analysis. Cholesterol and albumin levels from serum analysis **F)** Red and white blood cell counts from automated cell counter. Neutrophil and lymphocytes were manually counted from a prepared slide. **G)** H&E sections of liver, lung, and spleen from iEF treated mice.

We first evaluated the safety profile of iEF treatment in mice by analyzing changes in body weight, visceral organ histology, and whole blood counts. Immunocompetent and non-tumor bearing BALB/c mice were exposed to either iEF or sham (non-powered) coils for 30 days. We did not observe any significant changes in body weight between the iEF-treated and control groups (**Fig. 1D**). In addition, we did not observe any significant changes in behavior such as hunching and aggressive behavior in iEF-treated mice compared to control. Furthermore, no major significant differences were observed in whole blood count and blood biochemistry in iEF-treated mice compared to control (**Fig. 1E-F**). We also analyzed the histopathological features of vital organs like the liver, lungs, and spleen and observed no significant morphological and pathological changes in organs derived from iEF-treated mice compared to control (**Fig. 1G**). These results demonstrate that iEFs (as high as ∼1 mV/cm) can be safely applied to healthy mice without any adverse effects.

### iEFs inhibit tumor growth and lung metastases of TNBC

Next, we evaluated whether iEF treatment suppresses TNBC growth *in vivo*. We used an established orthotopic mouse breast tumor model where 4T1 cells were injected into the mammary fat pads of immunocompetent BALB/c mice (**Fig 2A**). Tumor bearing mice were placed in cages inside the iEF apparatus described earlier (**Fig. 2A**). Once tumors were palpable, mice were continuously exposed to iEF treatment for 15 days. We observed a significant difference in primary breast tumor volume and weight in iEF-treated mice compared to control (**Fig. 2B-C**). Moreover, iEF treatment of 4T1 cells *in vitro* showed significantly higher apoptosis compared to control (**Supplementary Fig. 1**). We also observed a significant difference in the number of lung metastases *in vivo* in iEF-treated mice compared to control by analyzing the histopathology of lung sections (**Fig. 2D-E**). In addition, we observed reduced maximum Feret diameter of metastases in lung H&E sections obtained from iEF-treated mice compared to control mice (**Fig. 2F**). The observations that iEF treatment reduced lung metastases prompted us to analyze the effect of iEF on epithelial-to-mesenchymal transition (EMT) in 4T1 cells *in vivo*. EMT plays a key role in breast cancer cell invasion and metastasis^21^. The hallmark of EMT is downregulation of epithelial markers such as E-cadherin and upregulation of mesenchymal marker such as N-cadherin^22^. Strikingly, we observed that iEF treatment significantly reduced the expression of N-cadherin and increased the expression of E-cadherin (**Fig. 2G**). These results suggest that iEF treatment limits metastatic outgrowth *in vivo* by inhibiting EMT in breast cancer cells.

**Figure 2:**
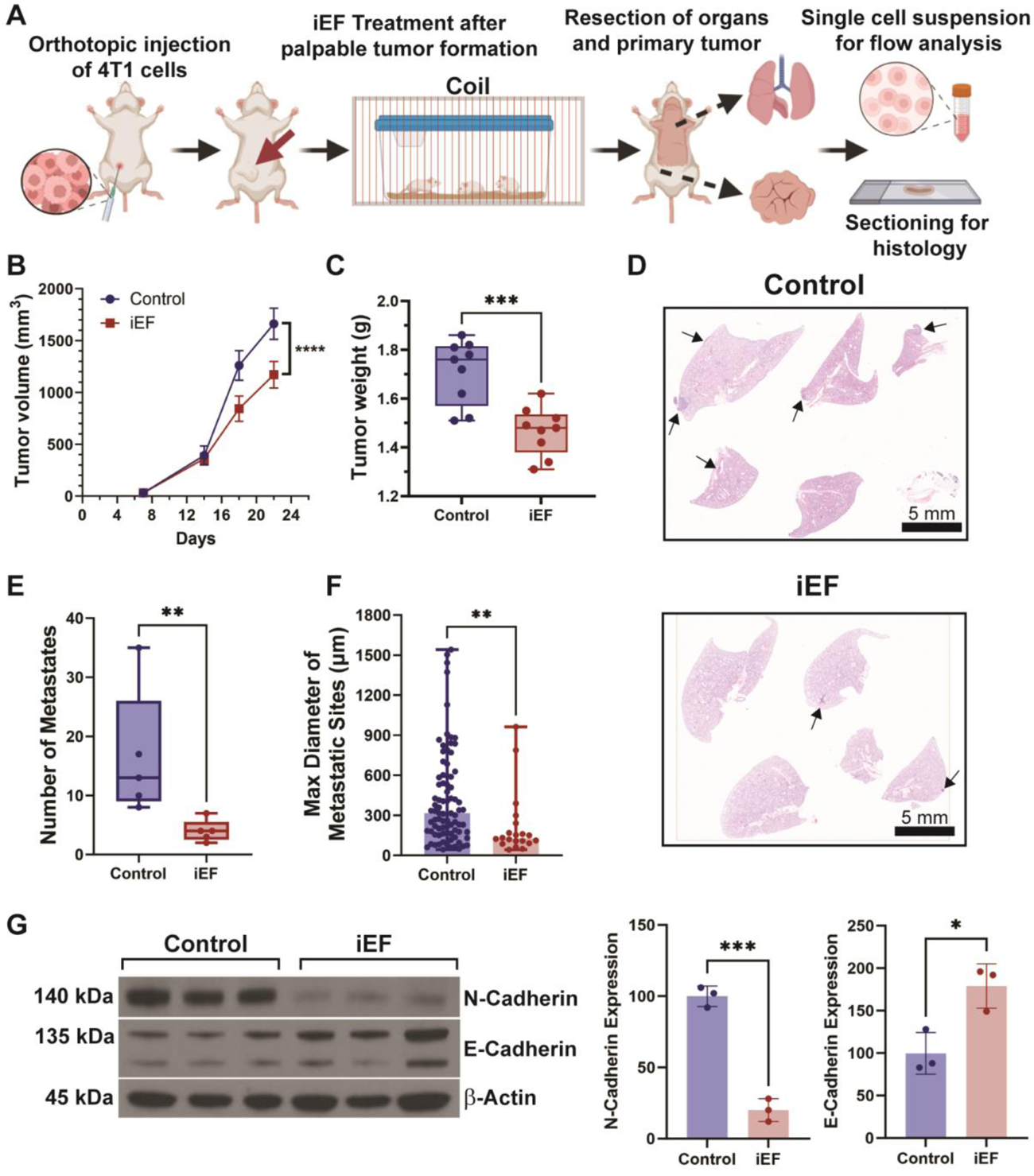
iEF inhibits TNBC tumor growth and metastasis. **A)** Graphical protocol for *in vivo* experiments. 4T1 cells were injected into the mammary fat pad of BALB/c mice. Treatment under electric fields began after palpable tumors had formed. Once the end point was reached organs and primary tumor were either digested into single cell suspension for flow cytometry or formalin fixed and sectioned for histology analysis. **B)** Tumor volume was measured over the course of iEF treatment and **C)** tumor weights were measured at the end of the experiment (n=9). **D)** Representative H&E-stained sections of mouse lungs harvested at the end of treatment. Black arrows indicate examples of metastases. **E)** Number of metastases identified in H&E sections of lungs (n=5). **F)** Maximum Feret diameter of metastatic sites in H&E sections of lungs. **G)** Western blots of EMT markers from primary tumor. * p < 0.05, ** p < 0.01, *** p < 0.001, **** p<0.0001.

Additionally, we analyzed serum samples collected at endpoint (day 15) from control or iEF treated 4T1-tumor bearing mice to determine the changes in the circulating cytokines due to iEF treatment. We observed a significant reduction in the serum levels of the cytokines IL-23 and CCL22 derived from the iEF-treated mice compared to control (**Fig. 3**). Sources for IL-23 and CCL22 in tumors include cancer cells and myeloid cells^23–26^. IL-23 is known to promote mammary tumor growth and lung metastasis, increase the infiltration of M2-like macrophages and neutrophils, and reduce the recruitment of CD8+ T cells in the TME^27^. CCL22 attracts Tregs to the breast tumor microenvironment and contributes to an immunosuppressive milieu^28,29^.

**Figure 3:**
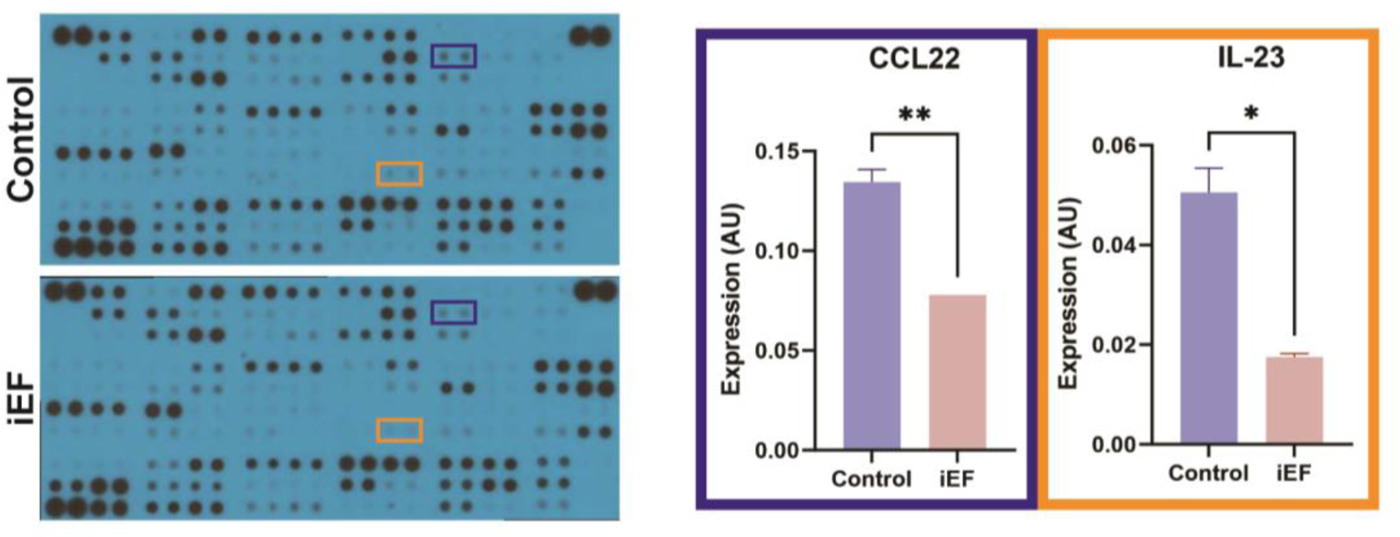
iEF treatment reduces circulation of pro-tumor cytokines. Blood serum was isolated from 4T1-tumor-bearing mice treated with iEFs or untreated control (as described in Fig. 2) and analyzed for cytokine and chemokine production by using a cytokine array kit. (*P < 0.05, **P < 0.01).

To confirm whether inhibition of breast tumor growth and metastasis by iEF treatment requires an intact immune system, we conducted experiments where 4T1 cells were implanted orthotopically into immunodeficient NOD *scid* gamma (NSG) mice. We observed no significant difference in 4T1 tumor growth and metastasis when grown in immunodeficient NSG mice (**Fig. 4**). Thus, when the results from the immunocompetent BALB/c (**Fig. 2**) and immunodeficient NSG (**Fig. 4**) mouse models are examined together, our findings suggest that iEF mediates anti-tumor and anti-metastatic effects by regulating the immune cells.

**Figure 4:**
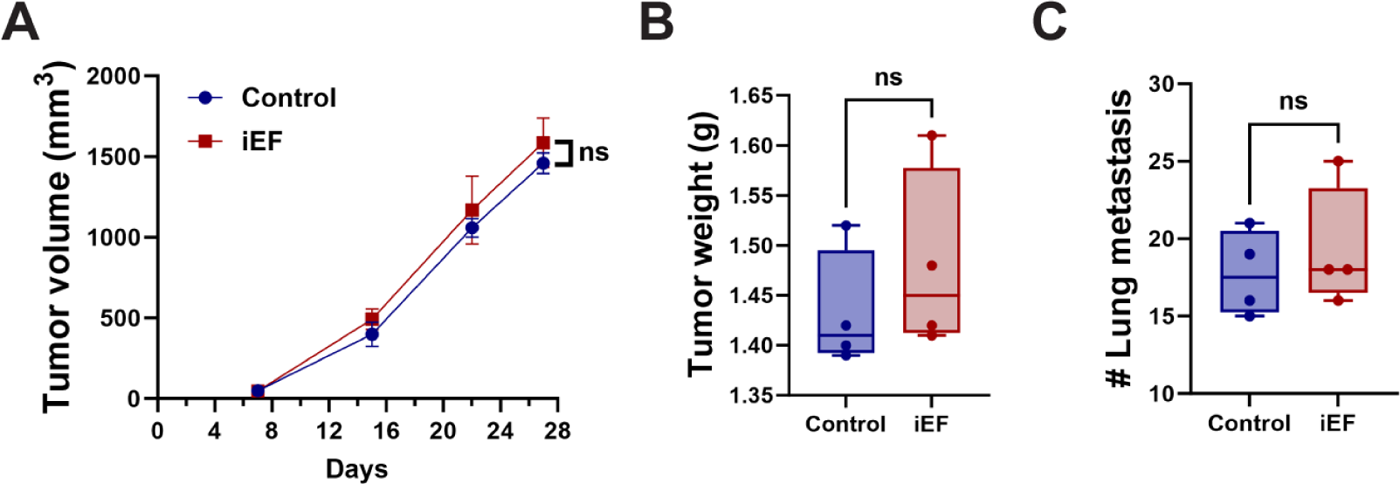
iEF treatment does not affect breast cancer growth and metastasis in immune-compromised mice. 4T1 cells were injected into the mammary fat pad of female NSG (NOD *scid* gamma) mice and treated with control or iEF (n=4). **A)** tumor growth was monitored every week and **B)** tumor weight was calculated at the end of the experiment. **C)** Lungs were harvested and metastasis was quantified.

### iEF treatment reduces immunosuppressive features of TNBC

The dynamic interplay between tumor and immune cells significantly influences the metastatic potential of tumor cells^30,31^. Our findings that suppression of tumor progression and metastasis by iEF treatment was contingent on an intact immune system led us to analyze the effect of iEFs on the tumor immune microenvironment. The migratory behavior of macrophages can influence tumor progression^32,33^ and plays a crucial role in coordinating immune responses and inflammation^34,35^. Thus, we first assessed the direct effect of iEF treatment on immune cell migration. Using an *in vitro* Transwell assay, we observed that RAW 264.7 (murine monocyte/macrophage-like cell line) cell migration was significantly reduced in iEF-treated groups compared to control (**Supplementary Fig. 2**). These results demonstrate that iEF treatment directly hinders immune cell migration.

Next, we analyzed the *in vivo* effects of iEF treatment on the modulation of immune cells in TNBC. Using the 4T1 orthotopic model as described earlier (**Fig. 2A**), we analyzed the single-cell suspensions of the tumor by high-dimensional multi-spectral flow cytometry. In a separate set of experiments, 4T1 tumors were treated with iEF for 8 days and primary tumors were harvested. We observed a significant decrease in the infiltration of regulatory T cells (Tregs) within the tumor in the iEF treatment group compared to control (**Fig. 5A**). Tregs are known for suppressing immune responses, and their reduction could indicate a shift towards a more anti-tumor immune environment^36^. In addition, iEF-treated tumors show decreased frequency of TIM3+ and PD-1+ CD8+ T cells (**Fig. 5B**). TIM3+PD-1+ co-expressing CD8+ T cells have been demonstrated to show increased exhaustion phenotype^37^. Furthermore, TIM3+PD-1+ CD8+ T cells from iEF-treated tumors show significantly reduced expression of CD39, an ectonucleotidase expressed by terminally exhausted T cells (**Fig. 5C**). CD8+ T cells play a crucial role in targeting and eliminating tumor cells, and a reduction in exhaustion features in CD8+ T cells may improve their functionality and enhance anti-tumor immune responses.

**Figure 5:**
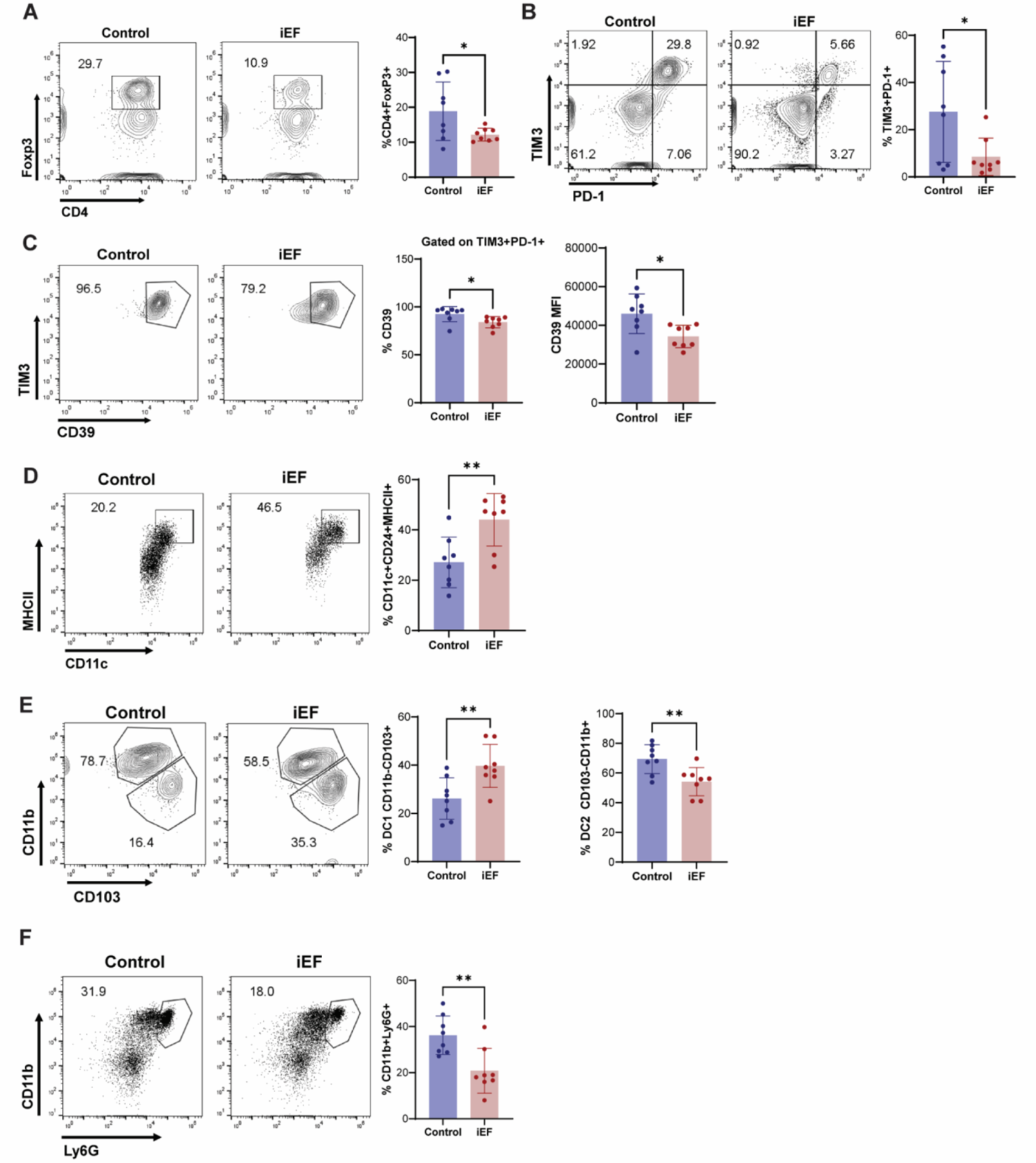
iEF treatment reduces immunosuppressive tumor microenvironment and promotes antitumor immunity in an orthotopic breast cancer model. 4T1 cells were injected in the mammary fat pad in female 6-to 8-week-old BALB/c mice and treated with iEF. Control or tumors treated with iEF were analyzed for the recruitment of various immune cells by flow cytometry. Representative flow cytometry plots for the tumors are illustrated. Flow cytometry data was used to analyze the levels of **A)** Tregs (CD4+Foxp3+), **B)** CD45+CD3+CD8+TIM3+PD-1+, **C)** CD45+CD3+CD8+TIM3+PD-1+CD39, **D)** CD45+CD11c+CD24+MHCII+, **E)** CD45+CD11b-CD103+ (DC1) and CD45+CD11b+CD103-(DC2), **F)** CD45+CD11b+Ly6G+ (granulocytic). p values were calculated by the student’s t-test. (*p < 0.05, **p < 0.01).

We also found that iEF treatment increased the abundance of dendritic cells (DCs) expressing elevated levels of major histocompatibility complex class II (MHCII) molecules (**Fig. 5D**). MHCII molecules are critical for presenting antigens to CD4+ T cells, and an upregulation suggests an enhanced capacity for the presentation of exogenously derived tumor-specific peptide antigens, potentially leading to a more robust anti-tumor immune response^38^. Additionally, iEF treatment increased the enrichment of Type 1 DCs (DC1s) compared to the control group (**Fig. 5E**). DC1 promotes Th1 immune responses and enhances cytotoxic T cell activity^39^, thereby indicating a positive shift towards an anti-tumor immune state. We further observed a reduction in Ly6G+ neutrophils in the iEF treatment group compared to the control group (**Fig. 5F**). However, we did not observe any significant changes in Ly6c+ monocytic cells (**Supplementary** Fig. 3). Neutrophils are known to possess immunosuppressive properties and have been linked to supporting tumor progression^40^. Collectively, our findings suggest that iEF treatment induces a multifaceted anti-tumor response by reducing the immunosuppressive features of the breast TME.

### iEF treatment regulates the immune infiltrates at the metastatic site in the lung

The unique microenvironment of distant organs may attract specific immune cell populations that facilitate metastatic colonization^41^. The observed inhibition of metastasis (**Fig. 2**) and immunomodulatory effects of iEFs in the primary breast TME (**Fig. 5**) prompted us to investigate the immune features of the metastatic lung microenvironment mediated by iEF treatment. We used the 4T1 orthotopic model as described earlier (**Fig. 2A**). 4T1 tumors were treated with iEF for 8 days and lungs were harvested. Single-cell suspensions of the lungs were analyzed by using a high-dimensional multi-spectral flow cytometry. We found that iEF-treated mice showed increased recruitment of CD8+ T cells at the metastatic site in the lungs (**Fig. 6A**). The CD8+ T cells also showed high Ki-67 expression (**Fig. 6B**), which suggests that the CD8+ T cells in the lung microenvironment are actively dividing and potentially exerting a more robust immune response^42^. In addition, iEF treatment increased the frequency of CX3CR1-expressing effector CD8+ T cells (**Fig. 6C**). However, we did not observe any significant changes in the frequency of TIM3 and PD-1 co-expressing exhausted CD8+ T cells (**Supplementary Fig. 4A**). Importantly, CD8+ T cells have been demonstrated to directly inhibit lung metastasis in TNBC^42^. Moreover, iEF treatment caused an increased infiltration of B cells (CD19+) in the lung microenvironment (**Fig. 6D**). B cells play an important role in adaptive immunity^43^. Of note, these CD19+ B cells in the lung microenvironment show reduced PD-L1 (programmed death ligand 1) expression (**Fig. 6E**). PD-L1 is known to be an immune checkpoint molecule, and reduced expression suggests a potentially more favorable immune environment^44^.

**Figure 6:**
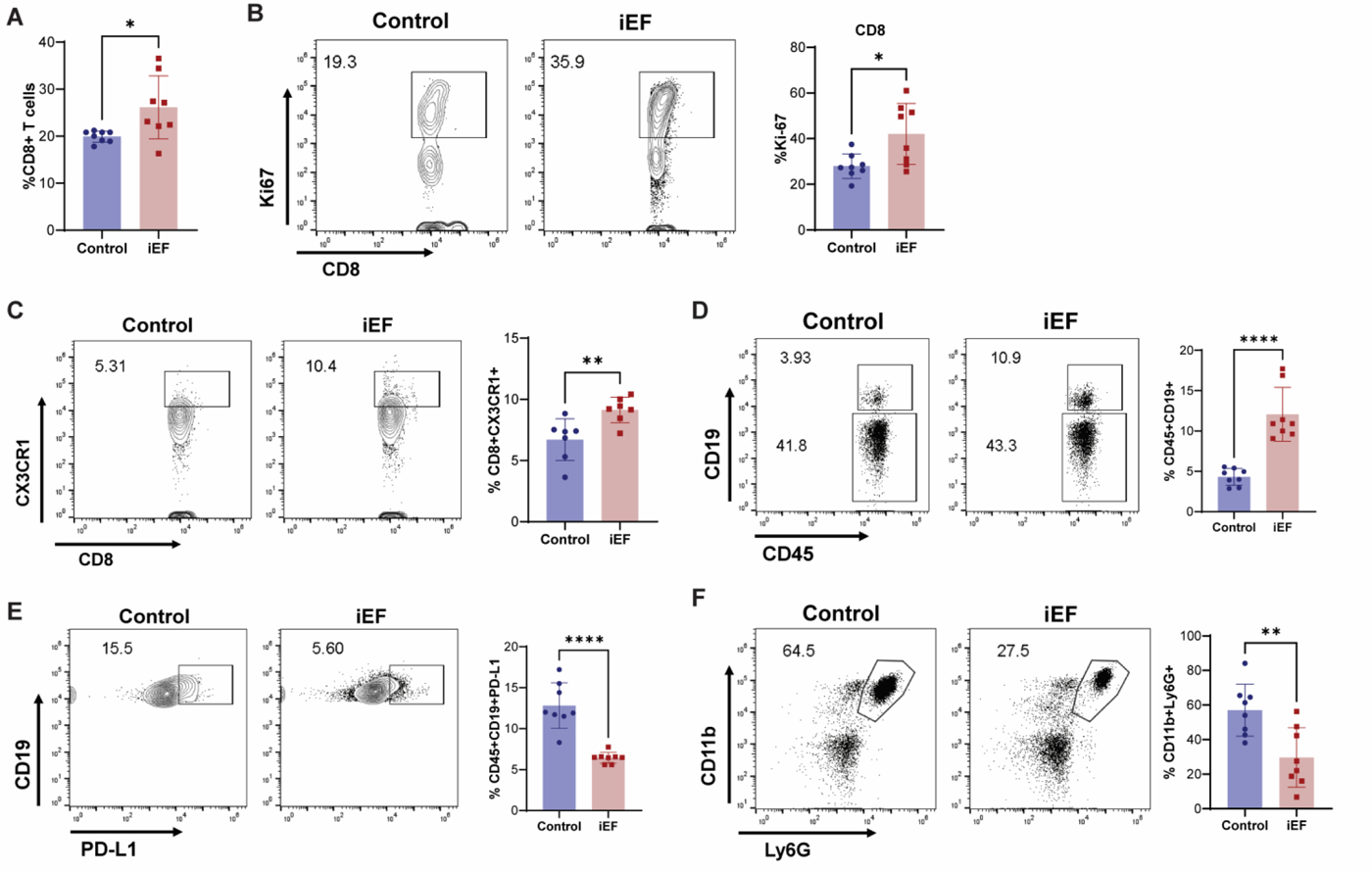
iEF regulates the immune cell recruitment to the metastatic site in the lung. 4T1 cells were injected in the mammary fat pad in female 6-to 8-week-old BALB/c mice and treated with control or iEF. Lungs from control or iEF-treated groups were analyzed for the immune infiltrates by flow cytometry. Representative flow cytometry plots for the tumors are illustrated. Flow cytometry results revealed the changes in **A)**CD45+CD3+CD8+, **B)** CD45+CD3+CD8+Ki-67+, **C)**CD8+CX3CR1+, **D)** CD45+CD19+, **E)** CD45+CD19+PD-L1+, **F)** CD45+CD11b+Ly6G+ (granulocytic). p values were calculated by the student’s t-test. (*P < 0.05, **P < 0.01, ****P < 0.0001).

iEF treatment also decreased the enrichment of CD11b+Ly6G+ neutrophils in the lung microenvironment (**Fig. 6F**). However, the Ly6c+ monocytic immune cells were unchanged (**Supplementary Fig. 4B**). Notably, CD11b+Ly6G+ immune cells are reported to inhibit CD8+ T cell functions and facilitate breast cancer metastasis^42^. Neutrophils also play a critical role in creating a pre-metastatic niche in the lung microenvironment and their reduction may contribute to a potentially less permissive environment for metastatic seeding and colonization in breast cancer^45,46^. The alterations in immune cell infiltrates, such as increased CD8+ T cells, B cells with reduced PD-L1 expression, and decreased neutrophils suggest that iEF treatment hinders metastasis by orchestrating changes in the immune landscape of the lung microenvironment.

## Discussion

We have shown that iEF technology can be a non-invasive approach to inhibit tumor growth and metastasis in a mouse TNBC model. To our knowledge, this is the first demonstration of an alternating electromagnetic, contact-free, and non-pharmacological means of inhibiting TNBC growth and lung metastasis *in vivo*. iEF treatment of tumor-bearing mice enables effective anti-tumor immune responses both in the primary TME and in the lung metastatic site. Furthermore, our results indicate that iEF treatment is highly safe and does not show any adverse effects on weight, blood count, and histopathological characteristics of visceral organs in normal tumor-free mice. Thus, our findings suggest that iEF technology is a relatively safe treatment modality with a favorable therapeutic index.

We observed that 4T1 TNBC cells undergo apoptosis when treated with iEFs *in vitro*. This finding is notable, especially in the context of bioelectricity, as other methods of applying electrical current or fields, such as tumor treating fields^47^ (TTFs) and positive electrostatic charge therapy^48,49^ (PECT), have also reported selective apoptosis of cancer cells. In addition, a recent study showed that TTFs enhance anti-tumor immune activation in a murine glioblastoma (GBM) model^50^. The bioelectric nature of the TME has been recently revealed clinically^51^, which may be attributed to the presence of intracellular and extracellular endogenous electric fields^52–57^. Moreover, differences in electrical properties in tumor tissue compared to normal tissue have been reported^51,58,59^. Thus, the intriguing outcome that three different techniques utilizing electric fields (iEFs, TTFs, and PECT) were independently shown to induce selective apoptosis of cancer cells suggest that the processes exploit intrinsic bioelectrical differences between cancer cells and normal cells.

Most studies on the effects of electric fields in cancer, including ones with iEFs, have been in *in vitro* systems and monocultures of cancer cells. Yet, the complex interactions between cancer cells, immune cells, and their microenvironment are pivotal for mediating tumor progression and metastasis^60^. A key finding from our study is that iEFs enhanced host immune responses at both the primary mammary tumor and the metastatic lung sites *in vivo* (**Fig. 7**). In mammary tumors, iEFs reduced the enrichment of immunosuppressive cells such as Tregs and Ly6G+ neutrophils. In addition, iEF treatment reduced the exhausted phenotype of anti-tumor CD8+ T cells. We also found reduced production of CCL22 with iEF treatment, which is an immunosuppressive cytokine and contributes to the recruitment of Tregs. Notably, iEF treatment increased the number of DCs, which are critical for cross-presenting antigens to CD8+ T cells for generating an efficient anti-tumor immune response. In addition, the apoptotic effect of iEFs on cancer cells may promote the release of tumor antigens, enhancing their uptake and cross-presentation by DCs to activate CD8+ T cell-mediated immune responses. Thus, our results suggest that iEF treatment in a TNBC tumor bearing mice alleviates the immunosuppressive state of the breast cancer microenvironment.

**Fig. 7:**
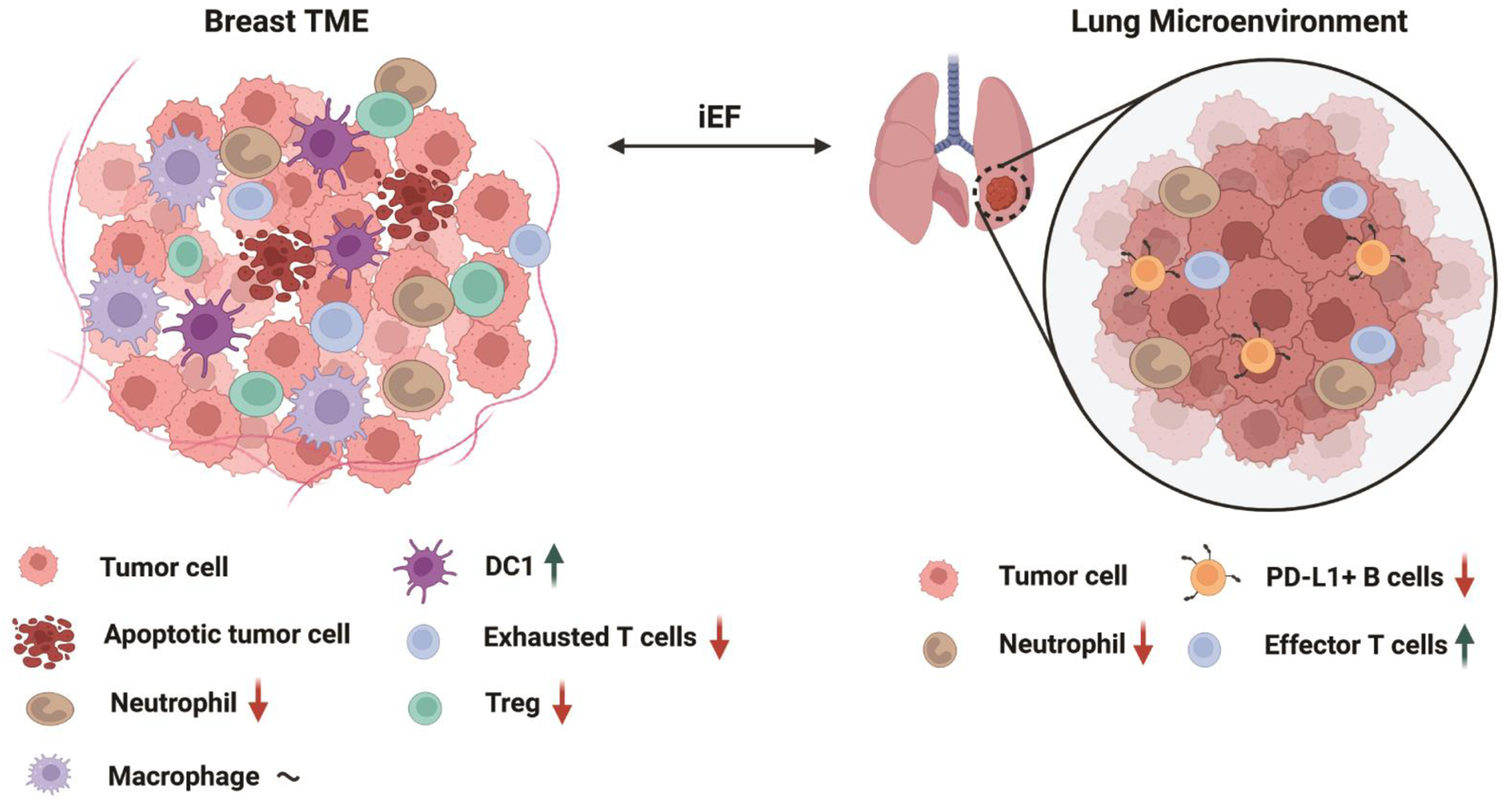
Immunomodulatory effects of iEF in the primary tumor and secondary metastatic site (lung).

In addition to the primary breast site, a robust immune response was observed at the metastatic site of the lungs due to iEF treatment. In the lungs, iEFs increased the population of proliferating (Ki-67 high) CD8+ T-cells, which suggests that the CD8+ T cells in the lung microenvironment are rapidly expanding and exerting a more robust immune response^42^. Studies have shown that CD8+ T cells play a direct inhibitory role in the metastasis of TNBC^42^. Concurrently, a decline in the pro-tumor CD11b+Gr1+ immune cell population within the pulmonary microenvironment was also observed, signifying a shift towards an anti-tumorigenic milieu. CD11b+Ly6G+ immune cells have been shown to suppress CD8+ T cell functions and thereby promote TNBC metastasis^42^. Furthermore, iEF treatment reduced the expression of PD-L1 in B cells in the lung microenvironment. PD-L1 is a ligand that binds to the PD-1 (programmed cell death protein 1) receptor on T cells, inhibiting T cell activation and proliferation^61^. PD-L1-expressing B cells may directly interact with PD-1-expressing T cells and inhibit their function thereby diminishing the anti-tumor response. This may contribute to immune tolerance within the lung microenvironment, allowing cancer cells to proliferate and evade immune surveillance. The enrichment of PD-L1+ B cells correlates with higher tumor grade in breast cancer^62^. In addition, melanoma patients show a higher frequency of PD-L1+ B cells at the metastatic site and suppress T cell activity in a PD-L1-dependent manner^63^. However, the mechanism of PD-L1 upregulation in B cells is not well understood.

To date, there is a dearth of studies on the effects of electric field treatment in *in vivo* cancer models. Here we demonstrate the feasibility of conducting long-term experiments on mice using a novel iEF apparatus. Yet, a limitation of the experimental design of the present study is the iEF apparatus surrounds a mouse cage and therefore subjects an entire mouse to continuous iEF treatment. Hence, we cannot discern the impact of iEF treatment locally to the tumor only versus systemic effects. The coils that generate iEFs can be embedded within a wearable garment to help enable treatment local to an implanted tumor. However, mice have the propensity to impart damage on any clothing worn, thereby rendering the electronics inside the garment non-functional. Thus, the configuration of the described iEF apparatus was important to ensure reproducible and uninterrupted iEF treatment of tumor-bearing mice.

In summary, we have developed a method for the successful application of iEFs *in vivo* in a non-contact and non-invasive manner. We have shown that iEF treatment inhibits metastatic outgrowth of breast cancer *in vivo* as well as inhibiting EMT. Strikingly, we observed that iEF treatment modulates an anti-tumor immune response at both the primary tumor and secondary metastatic site. To our knowledge, this is the first time that such a comprehensive and robust host immune response prompted by iEF treatment *in vivo* has been reported.. The novel findings from our study also point to the need for further investigation of the effects of alternating electric fields on the complex interactions that coordinate immune responses during cancer progression. Moreover, the immunomodulatory effects of iEF treatment may provide the foundation for harnessing anti-tumor immunity and expanding clinical responses to immunotherapy in the treatment of metastatic breast cancer and other solid tumors. This underscores the importance of exploring iEFs as a novel and bioelectric-based or electrotherapeutic intervention for cancer patients.

## Materials and Methods

### Cell culture

4T1 and RAW 264.7 cells were purchased from ATCC. 4T1 and RAW 264.7 cells were cultured in RPMI-1640 (**ATCC**) supplemented with 10% FBS (**Sigma-Aldrich**) and 1% penicillin/streptomycin (**Lonza**).

### Animal studies

All experiments were approved by the Institutional Animal Care and Use Committee of The Ohio State University (2007A0233-R5). Animals were housed according to University Laboratory Animal Resources guidelines. BALB/c (immuno-competent) and NOD *scid* gamma (NSG, immunodeficient) mice were purchased from Jackson Laboratory. Mice were injected with 4T1 cells (1x10^5^ cells in 100 µL PBS) directly into the fat pad of 4^th^ mammary gland. Tumor-bearing mice were randomly allocated into either iEF or sham treatment groups. Treatment began once palpable tumors had formed and continued until reaching endpoints. Palpable tumors were measured weekly using calipers. Tumor volume was calculated using the formula V = 4π/3 x (D_L_/2) x (D_S_/2)^2^ where D_L_ is the largest and D_S_ is the smallest superficial Feret diameter. After treatment, mice were euthanized, and primary tumors and lungs were harvested for further analysis.

### Application of iEF

For *in vivo* experiments, *once* tumors were palpable, mice were transferred to sterile cages with bottled water supply and the entire cage slid into the bore of the coil for treatment. The coil was constructed of ¼”-thick acrylic with outer dimensions of 20x21.5x39.5 cm (height x width x length). The housing was wrapped with 32 AWG magnetic wire with a total of 62 equally spaced turns (**Supplementary Fig. 5**). The DC resistance of the coil was 28 Ω. The inductance of the coil was measured by an LCR meter (**Keysight U1733C**) to be 437 μH at 100 kHz. The applied fields were generated by applying a 100kHz, 20Vpp sawtooth waveform (**Hewlett Packard 33120A**) which generated a peak magnetic and electric field strength of 5.5 µT and 1.2 mV/cm respectively in the plane where the mice are free to move.

While the iEF properties cannot be directly measured *in vivo*, they can be calculated from Maxwell’s equations (see **Supplementary**). The calculated iEF properties were validated (see **Supplementary**) experimentally by comparing calculated values of the magnetic induction and comparing with point-wise measurements of inside the box using a flux gate magnetic sensor (**Magnetic Sciences, Model#MC162**). The variation of field strength throughout the cage can be seen in **Figure 1B** and **Supplementary Fig. 6**.

For *in vitro* experiments, smaller, similarly constructed coil was used. The coil was constructed with acrylic with outer dimensions of 8 x 9.5 x 13.5 cm (height x width x length). The housing of the coil was wrapped with X equally spaced turns of 32 AWG magnetic wire (**Supplementary Fig. 7**). The DC resistance of the coil was 4.2 Ω and the inductance of the coil was measured by an LCR meter (**Keysight U1733C**) to be 27 µH at 100 kHz. The applied fields were generated by applying a 100kHz, 20Vpp sawtooth waveform (**Hewlett Packard 33120A**) which generated a peak magnetic and electric field strength of 25 µT and 3.2 mV/cm. The variation of the calculated magnetic and electric field strength can be seen in **Supplementary** Fig. 8.

### Flow cytometry

Mouse tumor or lung samples were mechanically disrupted and subjected to digestion with collagenase-IV (1.5 mg/ml; **Gibco**) for 45 min at 37°C with intermittent vortexing. The digested tissues were subsequently passed through 70 μm filters and layered on a discontinuous Percoll gradient (**Cytiva**). Centrifugation was carried out at 1800 rpm for 20 minutes without braking, resulting in the collection of immune cells at the gradient interface. These isolated cells were stained at 4°C with Live/Dead Blue viability dye for 10 minutes (**Invitrogen**) to exclude the nonviable cells. The cells were stained for extracellular surface markers for 30 minutes. All intracellular staining was performed using the Foxp3 TF staining kit (**Invitrogen**) according to the manufacturer’s instructions. All samples were recorded on a Cytek Aurora high-dimensional flow cytometer. All fluorochrome-conjugated primary antibodies used for flow cytometry are provided in **Supplementary Table 1**. The gating strategy for isolating cell populations is shown in **Supplementary Figure 9**. The data was analyzed using FlowJo software.

### Histopathology

Lung sections were mounted on microscope slides and digitally scanned. Metastatic sites were counted manually by visual inspection with a minimum diameter of 40 µm constituting a metastatic site.

### Immunoblotting

Immunoblotting was performed following the method as described^64^. Briefly, cell lysates were electrophoresed on NuPAGE 4–12% gradient precast gels (**Invitrogen**), and subsequently transferred onto nitrocellulose membranes with a pore size of 0.45 μm (**BioRad**). These membranes were then incubated with primary antibodies at specific dilutions (**Supplementary Table 2**). We used HRP-conjugated secondary antibodies, specifically goat anti-rabbit IgG for detection, and a chemiluminescent substrate (**Millipore**) was used for protein visualization. β-actin was used for loading controls for the blots.

### Cytokine Array

The cytokine array kit was used as per the manufacturer’s protocol (**R&D Systems, ARY028**). Blood was drawn from mice (n=5) at endpoint (day 15) of iEF treatment and pooled together for analysis.

### Transwell Migration Assay

Transwell permeable membranes with 8 µm pore size (**Corning, CLS3422**) were coated with 80 µL of 10µg mL^-1^ fibronectin solution (in 1X PBS, **Sigma FC010**) and left to dry overnight at room temperature. RAW 264.7 cells were starved in 0.1% FBS media for 12 hr prior to the start of the assay. The cells were collected and resuspended at a concentration of 1 x 10^6^ cells mL^-1^ in 0.1% FBS media. The top chamber of each Transwell was filled with 100 µL of cell suspension while the bottom chamber was filled with 650 µL of either 0.1% FBS media or CSF-1 supplemented media (50 ng mL^-1^ CSF-1). iEF treated plates were slid into a small rectangular coil (**Supplementary Fig. 7**) while control plates were slid into an unpowered identical coil (sham). Cells were allowed to migrate over 24 hrs. Afterward, cells were fixed and stained using a standard HEMA 3 solution kit (**Fisher Scientific, 23-123869**) and imaged using a BioTek Lionheart FX Automated Microscope. Migration values were quantified by counting cells in 5 separate fields of view on each membrane and normalizing by the number of seeded cells.

### Apoptosis Assay

4T1 cells were plated in 6-well culture plates in full growth media (10% FBS) at a concentration of 2x10^5^ cells per well. iEF treated plates were slid into a small rectangular coil (**Supplementary Fig. 7**) while control plates were slid into an unpowered identical coil (sham). Cells were allowed to grow for 48 hrs with or without iEF treatment. After, cells were detached using 0.05% Trypsin and combined with the supernatant which contained floating cells. The cells were then washed with cold PBS and stained using the FITC Annexin V Apoptosis Detection Kit I (**BD, RRID: AB_2869082**). Flow analysis was performed on a BD LSRFortessa.

### Statistical Analysis

Statistical analyses were performed with Prism software (**GraphPad Software Inc.**). The data is reported as meanL±Lstandard deviation. Significance was calculated using unpaired Student’s t-tests to assess the differences and associated P-values between various groups of data. Statistical significance was noted in the figures as *PL<L0.05, **PL<L0.01, ***PL<L0.001, ****PL<L0.0001 and ns: non-significant.

## Acknowledgments

This work was supported by an Idea Grant from Pelotonia and the Ohio State University Comprehensive Cancer Center (OSUCCC), an Accelerator Award supported by the Keenan Center for Entrepreneurship at Ohio State University and the Ohio Third Frontier Technology Validation and Start-up Fund, and the Cancer Biology from of the OSUCCC (P30CA016058). This work was also supported in part by the Mary Wieczynski Furnivall Cancer Research Fund.

## Contributions

M.C., T.H.J., D.K.A., N.A., V.V.S., R.K.G., and J.W.S. conceived and designed the experiments. M.C., T.H.J., and D.K.A. performed the experiments. M.C. & T.H.J. performed data analysis. The manuscript was written by M.C., T.H.J., N.A., V.V.S., R.K.G., and J.W.S.

