## Supplement for "Induced electric fields inhibit breast cancer growth and metastasis by modulating the immune tumor microenvironment"

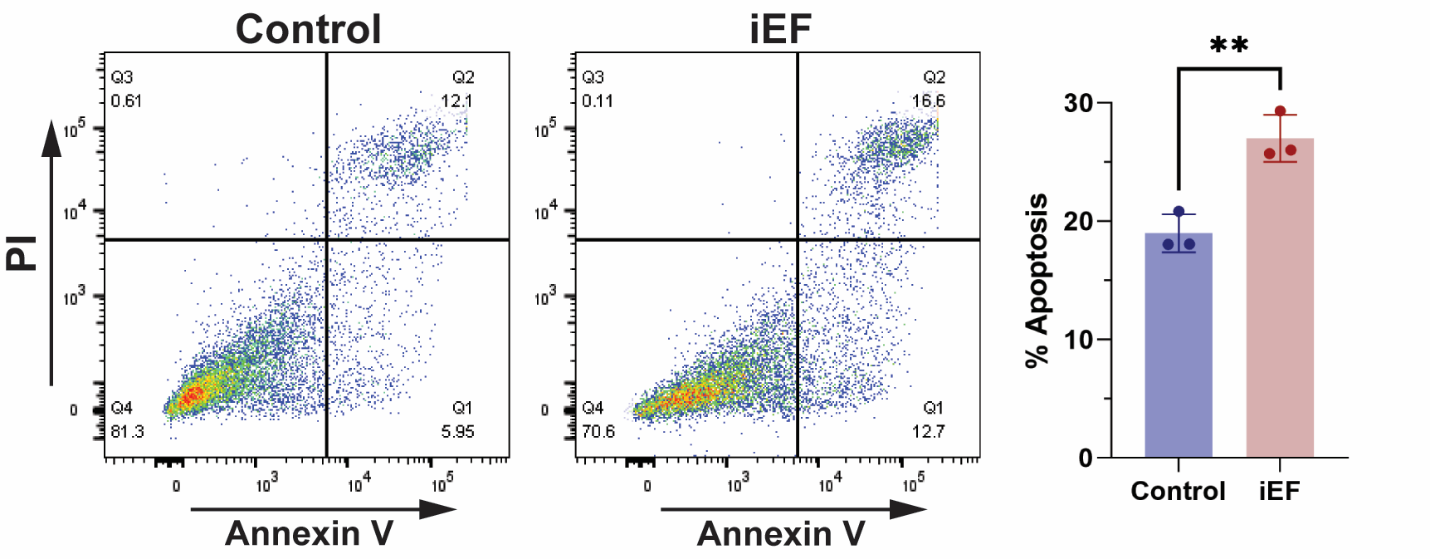

**Supplementary Figure 1:** Representative flow cytometry scatter plots of 4T1 cells treated with (iEF) or without (Control) iEF for 48 hours and stained using an Annexin V (FITC) and Propidium Iodide (PI) apoptosis kit. Apoptotic cells are those that are Annexin V positive (i.e. Q1 and Q2).

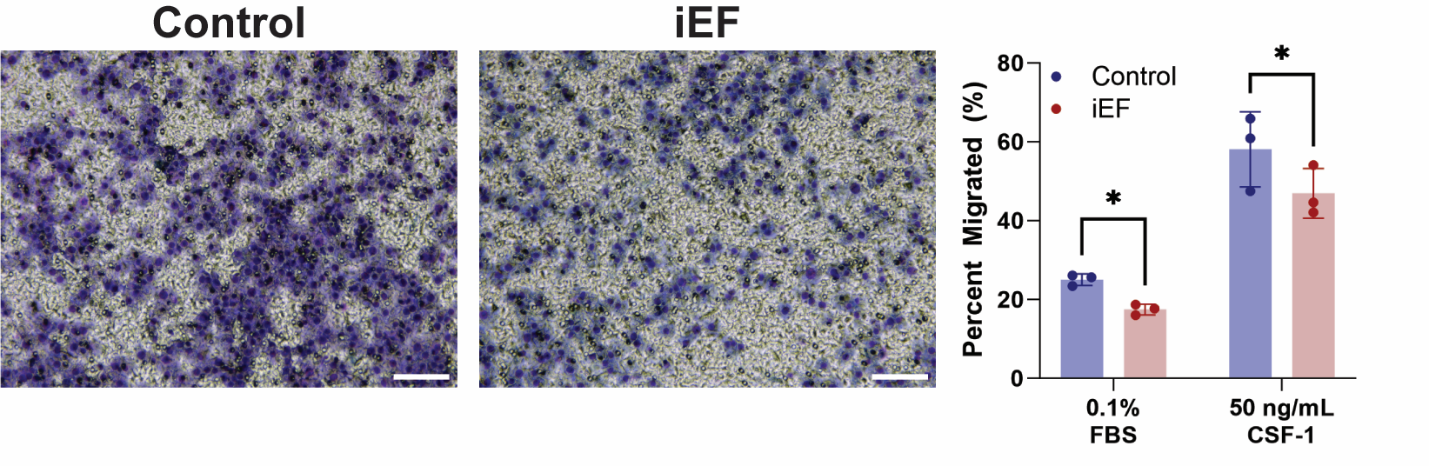

**Supplementary Figure 2**: Representative images of RAW 264.7 cells after 24 hrs of migration with 50 ng/mL CSF-1 as a chemoattractant in presence or absence of iEF in 8 µm Transwell inserts. Quantification of the migrated cells under different conditions in presence or absence of iEF. Scale bars are 100 µm.

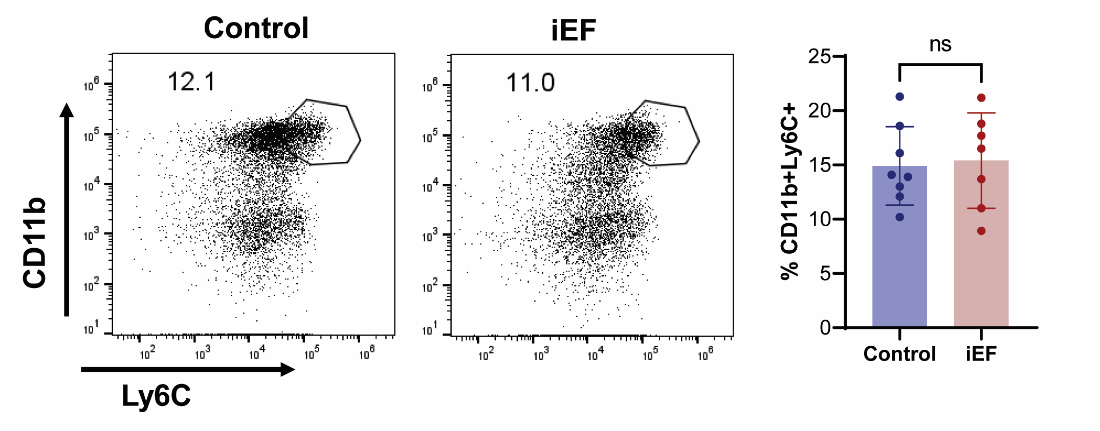

**Supplementary Figure 3**: CD45+CD11b+Ly6C+ (monocytic) cells from primary tumor.

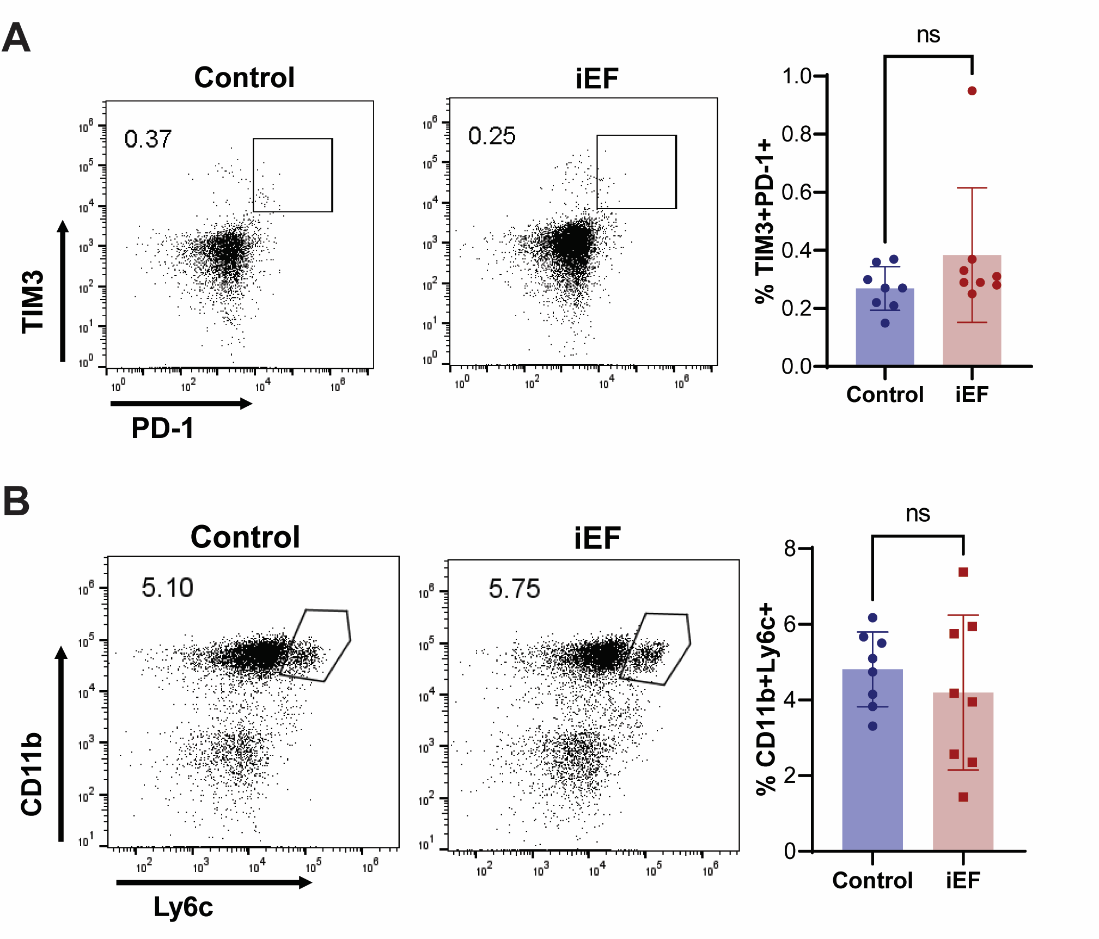

**Supplementary Figure 4**: **A)** CD45+CD3+CD8+TIM3+PD-1+ **B)** CD45+CD11b+Ly6C+ (monocytic) cells from lungs.

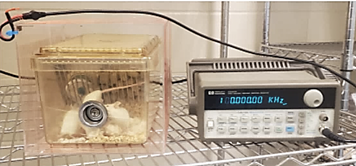

**Supplementary Figure 5**: Coil and generator used for mouse studies.

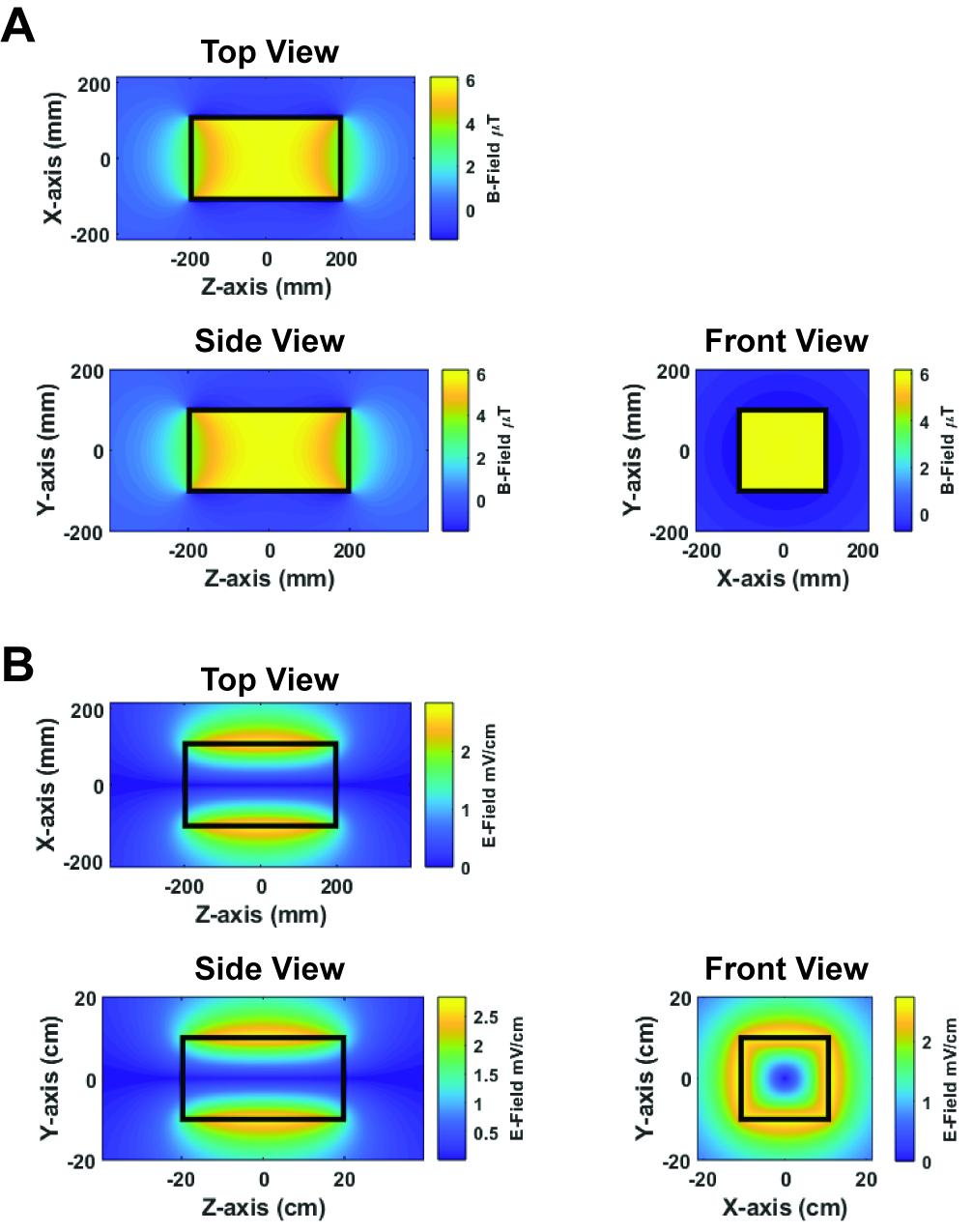

**Supplementary Figure 6**: **A)** Peak magnetic field intensity contours of mouse cage coil when driven by a 20 Vpp, 100kHz sawtooth waveform. Top view (top left), side view (bottom left), front view (bottom right). **B)** Peak electric field intensity contours of mouse cage coil when driven by a 20 Vpp, 100kHz sawtooth waveform. Top view (top left), side view (bottom left), front view (bottom right).

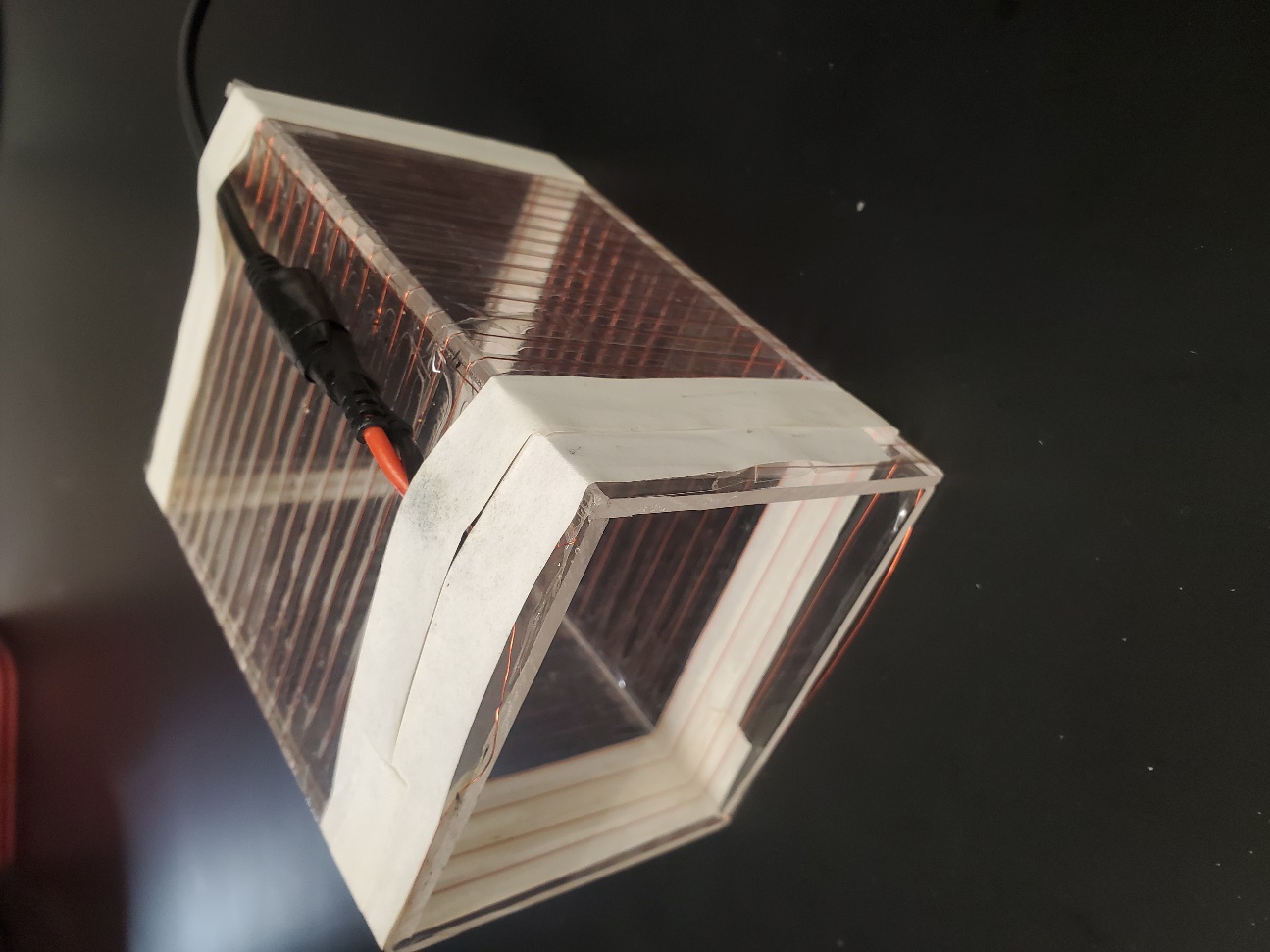

**Supplementary Figure 7**: In vitro Coil Image

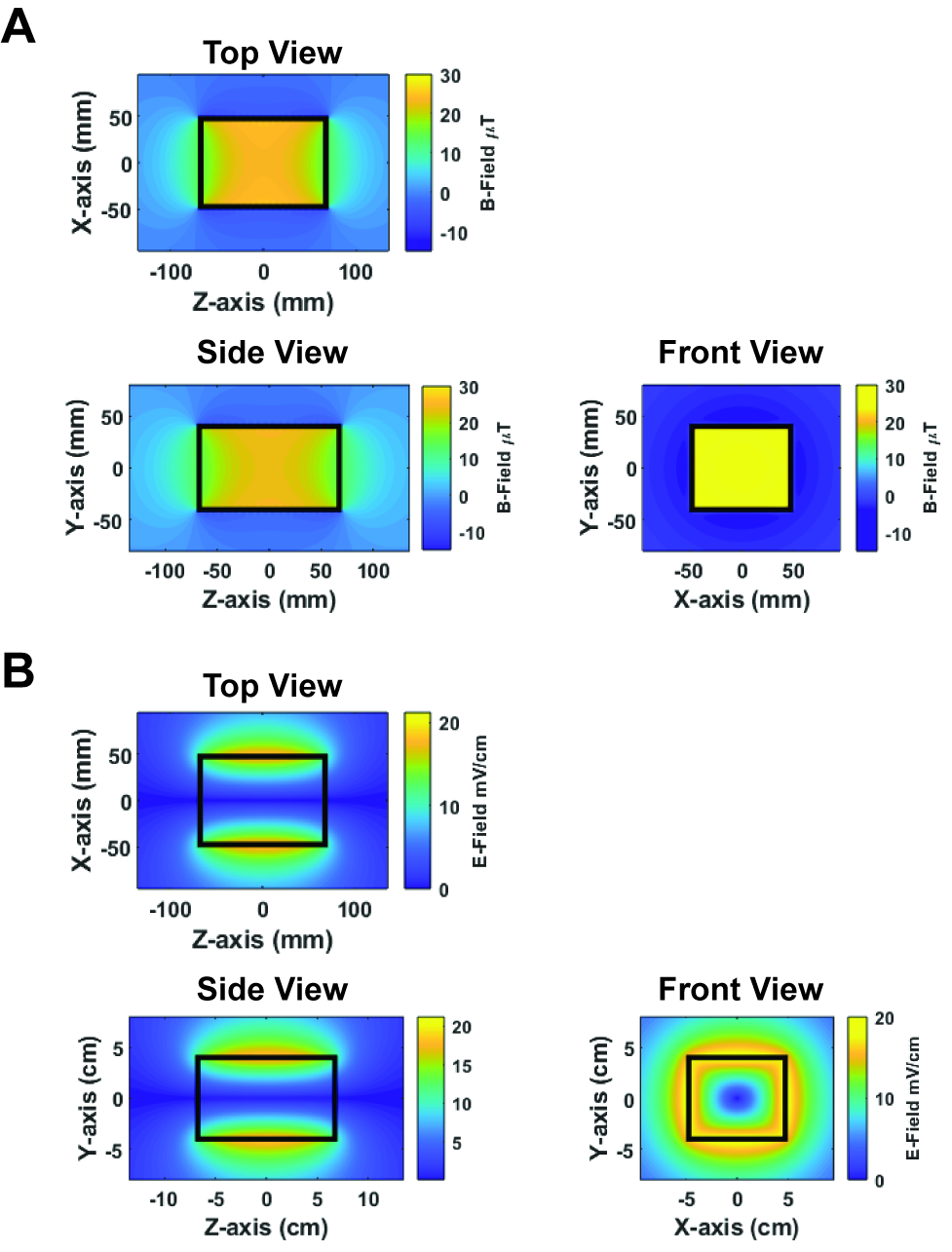

**Supplementary Figure 8**: **A)** Peak magnetic field intensity contours of *in vitro* coil when driven by a 20 Vpp, 100kHz sawtooth waveform. Top view (top left), side view (bottom left), front view (bottom right). (B) Electric field intensity contours of *in vitro* coil when driven by a 20 Vpp, 100kHz sawtooth waveform. Top view (top left), side view (bottom left), front view (bottom right).

**Supplementary Table 1**

List of flow cytometry antibodies:

| **Antibody** | **Fluorophore** | **Source** | **Catalog Number** |
| --- | --- | --- | --- |
| CD24 | BV480 | BD Biosciences | 752768 |
| CD3 | BUV737 | BD Biosciences | 612771 |
| CD39 | PerCP-Cy5.5 | BD Biosciences | 567270 |
| Ki-67 | BUV395 | BD Biosciences | 564071 |
| MHCll | BUV615 | BD Biosciences | 751570 |
| CD103 | BV711 | BioLegend | 121435 |
| CD11c | BV750 | BioLegend | 117357 |
| CD19 | Spark NIR 685 | BioLegend | 115567 |
| CD4 | APC Fire810 | BioLegend | 100479 |
| CD45 | BV510 | BioLegend | 103137 |
| CX3CR1 | APC/cy7 | BioLegend | 149047 |
| Ly6C | BV605 | BioLegend | 128035 |
| PD-1 | FITC | BioLegend | 135213 |
| PD-L1 | BV421 | BioLegend | 124315 |
| TCR β | PE/Cyanine5 | BioLegend | 109209 |
| Tim3 | BV711 | BioLegend | 134021 |
| CD11b | AF532 | eBiosciences | 58-0112-82 |
| FoxP3 | eflour450 | eBiosciences | 48-5773-82 |
| Ly6G | Super Bright 436 | eBiosciences | 62-9668-82 |
| Viability dye | Live Dead Blue | Invitrogen | L23105 |

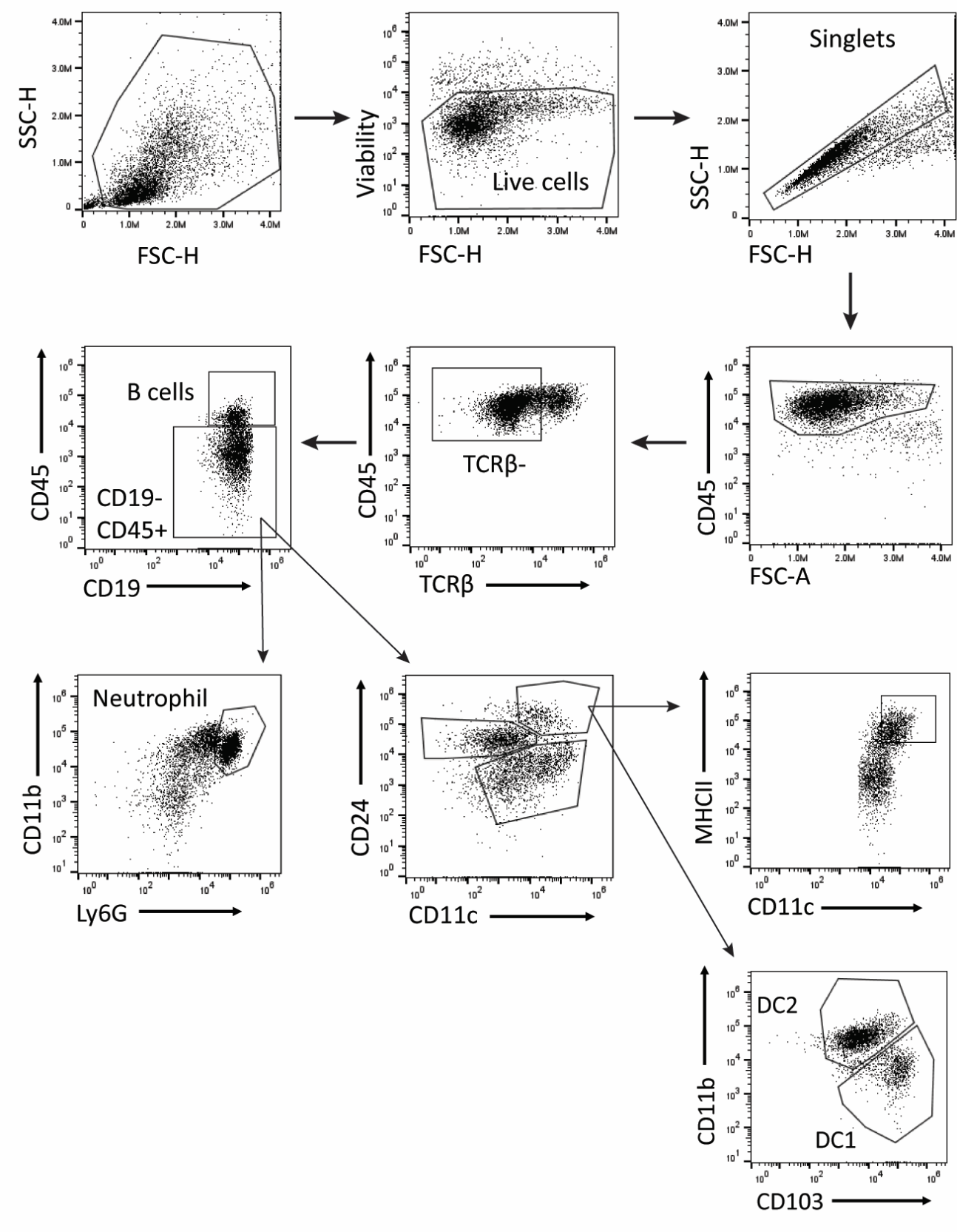

**Supplementary Figure 9**: Gating strategy of immune flow panel.

**Supplementary Table 2**

List of western blot antibodies:

| **Antibody** | **Source** | **Catalog Number** |
| --- | --- | --- |
| N-Cadherin | Cell Signaling Technologies | 13116 |
| E-Cadherin | Cell Signaling Technologies | 3195 |
| Β-Actin | Cell Signaling Technologies | 4970 |

**Field Calculation for Rectangular Coil**

For a coil of rectangular cross section, the equation for the components of the vector potential at any point in space is derived from the equations given in [1]:

$A_{x}\left( x,y,z \right)=\frac{\mu_{0}I(t)}{4\pi}\sum_{j=1}^{N} \sum_{k=1}^{2} \left[ \sinh^{-1} \left( \frac{x-a_{j}\left( -1 \right)^{1+k}}{r_{yz}} \right)-\sinh^{-1} \left( \frac{x-a_{j}\left( -1 \right)^{k}}{r_{yz}} \right) \right]$	**(1)**

$A_{y}\left( x,y,z \right)=\frac{\mu_{0}I(t)}{4\pi}\sum_{j=1}^{N} \sum_{k=1}^{2} \left[ \sinh^{-1} \left( \frac{y-b_{j}\left( -1 \right)^{1+k}}{r_{xz}} \right)-\sinh^{-1} \left( \frac{y-b_{j}\left( -1 \right)^{k}}{r_{xz}} \right) \right]$	**(2)**

where:

$$r_{yz}=\sqrt{\left( y-b_{j}\left( -1 \right)^{1+k} \right)^{2}+\left( z-c_{j} \right)^{2}}$$

$$r_{xz}=\sqrt{\left( x-a_{j}\left( -1 \right)^{1+k} \right)^{2}+\left( z-c_{j} \right)^{2}}$$

The induced electric field is given by:

$\mathbf{E}=-\frac{\partial\mathbf{A}}{\partial t}$	**(3)**

Substituting in **Equation 1** & 2:

$E_{x}=-\frac{dI}{dt}\frac{\mu_{0}}{4\pi}\sum_{j=1}^{N} \sum_{k=1}^{2} \left[ \sinh^{-1} \left( \frac{x-a_{j}\left( -1 \right)^{1+k}}{r_{yz}} \right)-\sinh^{-1} \left( \frac{x-a_{j}\left( -1 \right)^{k}}{r_{yz}} \right) \right]$	**(4)**

$E_{y}=-\frac{dI}{dt}\frac{\mu_{0}}{4\pi}\sum_{j=1}^{N} \sum_{k=1}^{2} \left[ \sinh^{-1} \left( \frac{y-b_{j}\left( -1 \right)^{1+k}}{r_{xz}} \right)-\sinh^{-1} \left( \frac{y-b_{j}\left( -1 \right)^{k}}{r_{xz}} \right) \right]$	**(5)**

Using **Equations 1** & **2**, the magnetic field is given by:

$B_{z}=\frac{\partial A_{y}}{\partial x}-\frac{\partial A_{x}}{\partial y}$	**(6)**

$=\frac{\mu_{0}I}{4\pi}\sum_{j=1}^{N} \sum_{k=1}^{2} \left[ \frac{x-a_{j}\left( -1 \right)^{1+k}}{r_{xz}^{2}}\left( \frac{y-b_{j}\left( -1 \right)^{k}}{\sqrt{\left( y-b_{j}\left( -1 \right)^{k} \right)^{2}+r_{xz}^{2}}}-\frac{y-b_{j}\left( -1 \right)^{1+k}}{\sqrt{\left( y-b_{j}\left( -1 \right)^{1+k} \right)^{2}+r_{xz}^{2}}} \right)+\frac{y-b_{j}\left( -1 \right)^{1+k}}{r_{yz}^{2}}\left( \frac{x-a_{j}\left( -1 \right)^{k}}{\sqrt{\left( x-a_{j}\left( -1 \right)^{k} \right)^{2}+r_{yz}^{2}}}-\frac{x-a_{j}\left( -1 \right)^{1+k}}{\sqrt{\left( x-a_{j}\left( -1 \right)^{1+k} \right)^{2}+r_{yz}^{2}}} \right) \right]$	**(7)**

**Code Validation**

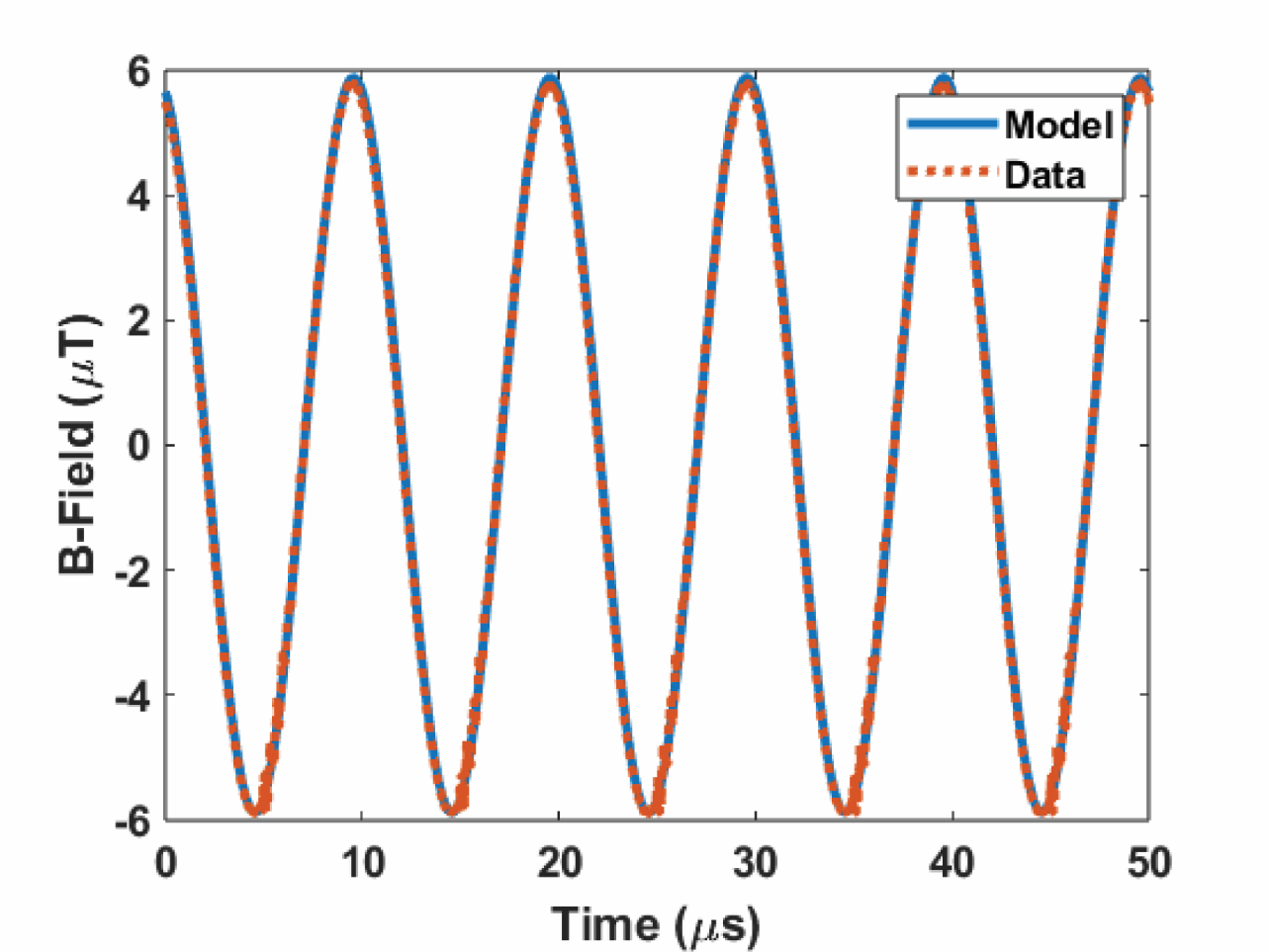

**Supplementary Figure 10:** Comparison of predicted and measured magnetic flux density (B-field) for the coil used in *in vivo* experiments. Field value is at center of coil when driven by a 20Vpp, 100kHz sine waveform.

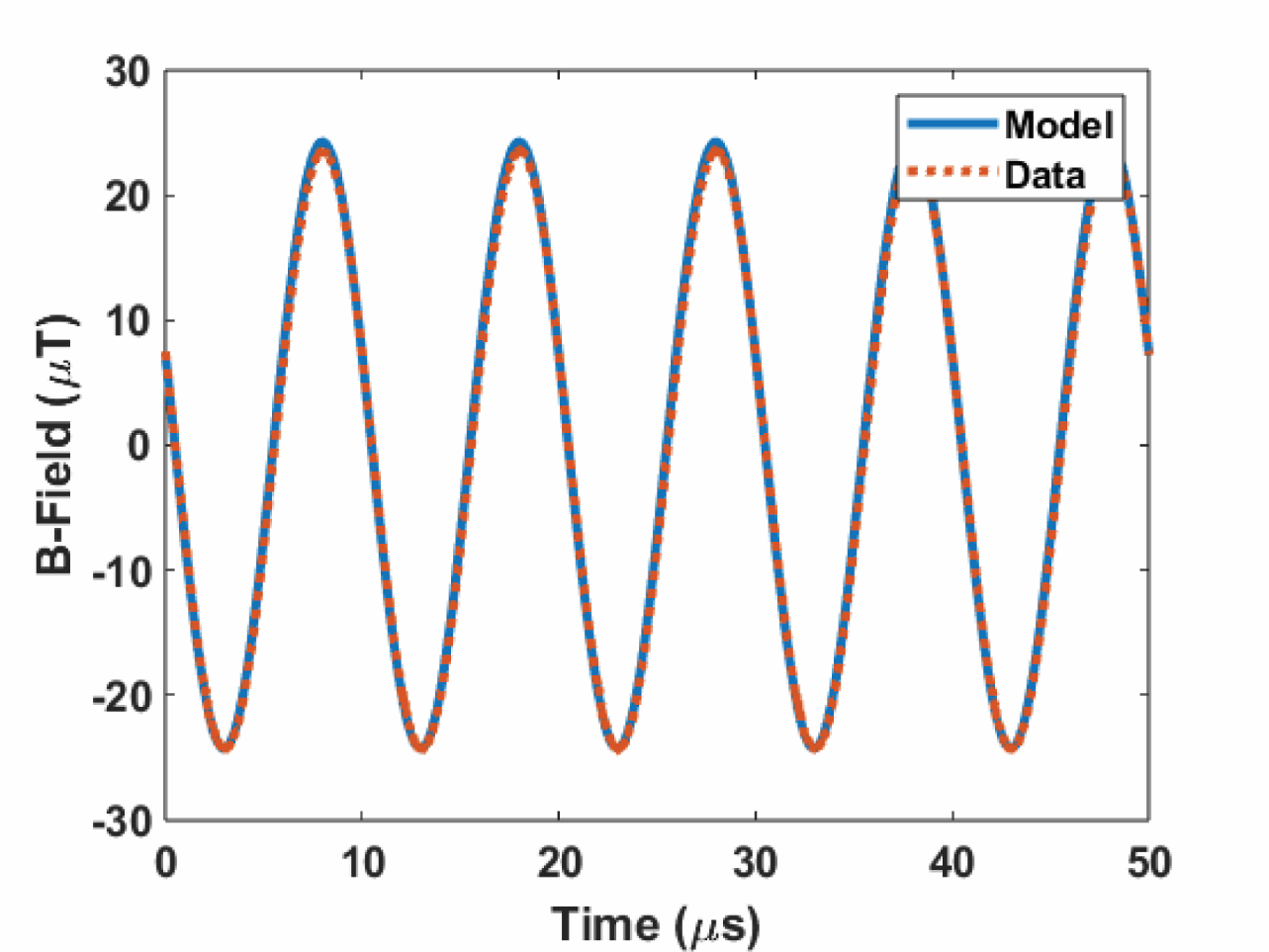

**Supplementary Figure 11:** Comparison of predicted and measured magnetic flux density (B-field) for the coil used in *in vitro* experiments. Field value is at center of coil when driven by a 20Vpp, 100kHz sine waveform.
